# Mechanical History and Substrate Stiffness Shape Integrin-Mediated Endothelial Cell Behavior on Bioactive Hydrogels

**DOI:** 10.64898/2026.07.28.741323

**Authors:** Abbey Nkansah, Aashee Budhwani, Ashauntee Fairley, Aarushi Chowdary Yedalla, Adhwita Anand, Nicholas Grammer, Josephine Allen, Elizabeth Cosgriff-Hernandez

**Affiliations:** Department of Biomedical Engineering, The University of Texas at Austin, Austin, Texas, 78712, USA; Department of Pediatrics, Medical College of Wisconsin, Milwaukee, WI, 53226, USA

**Keywords:** Integrins, Endothelial Cells, Bioactive Hydrogels

## Abstract

Synthetic blood contacting devices frequently fail due to the lack of requisite biochemical and biomechanical cues needed to support transanastamotic endothelialization. During transanastomotic endothelialization, endothelial cells experience dynamic changes in extracellular mechanical cues as they migrate from compliant native vessels onto stiffer blood contacting device surfaces. However, how substrate stiffness and mechanical memory from prior mechanical environments influence temporal integrin remodeling and downstream endothelialization processes necessary to establish a stable endothelial layer remains poorly understood. In this study, human coronary artery endothelial cells (HCECs) were cultured on substrates spanning physiologically relevant stiffnesses to determine how substrate mechanics regulate collagen binding integrins and endothelialization. Increasing substrate stiffness promoted time dependent upregulation of α2β1 integrin expression, whereas α1β1 expression remained unchanged. Enhanced α2β1 expression on stiff substrates was accompanied by increased vinculin associated focal adhesion maturation and accelerated endothelialization, characterized by increased proliferation, migration, and progression to confluence prior to reaching quiescence after 1 week. To better model transanastomotic migration and investigate mechanical history effects, cells initially expanded on compliant hydrogels were transferred to stiff substrates. Although these cells exhibited transient reductions in α2β1 expression at early timepoints compared with tissue culture polystyrene expanded controls, no persistent differences in focal adhesion maturation, proliferation, migration, confluence, or quiescence were observed. Collectively, these findings demonstrate that substrate stiffness is a primary regulator of the early endothelialization processes required to establish a stable endothelial monolayer, whereas the influence of mechanical history is transient and ultimately superseded by the current mechanical environment. These findings also identify α2β1 mediated mechanotransduction as a potential design target for blood contacting biomaterials that promote rapid endothelialization while supporting long-term endothelial cell quiescence.

## Introduction

Long-term thromboresistance remains a major limitation of synthetic blood-contacting devices, as these materials lack intrinsic biological mechanisms required to regulate hemostasis.^1–3^ In native vessels, endothelial cells maintain vascular hemostasis by forming a quiescent monolayer that actively suppresses thrombosis through tightly regulated hemostatic signaling.^4^ Consequently, the design of synthetic grafts has increasingly focused on recapitulating this native functionality by promoting endothelialization at the material interface.^5, 6^ Endothelialization is dictated by cell-material interactions that govern critical cellular functions such as adhesion, proliferation, and migration.^7–11^ These cellular responses facilitate the establishment of a quiescent phenotype.^4^ In addition to cell coverage, the endothelial layer must exhibit appropriate hemostatic function to establish long-term thromboresistance.^11, 12^

Central to these processes are integrin-mediated interactions with extracellular matrix ligands or adsorbed proteins, which transduce biochemical and mechanical signals that drive downstream phenotypic outcomes.^11–13^ In synthetic vascular grafts, the material surface chemistry largely determines the composition and extent of the adsorbed protein layer, which ultimately governs subsequent biological interactions.^14, 15^ The native basement membrane presents more defined ligand environments to promote endothelial cell attachment and hemostatic phenotype.^4, 11, 16–18^ As a result, strategies to improve graft performance in synthetic materials have focused on mimicking the native basement membrane through approaches such as extracellular matrix protein coatings or incorporation of bioactive ligands within engineered materials.^14, 19–22^ Critical to this strategy is understanding the interplay of integrin-mediated endothelial cell coverage.^2, 23^

As one of the key mechanisms for endothelialization on blood-contacting devices is transanastamotic migration, understanding the dynamics behind integrin-ligand interactions and biomechanical cues is critical. ^24, 25^ ^25, 26^ During transanastamotic migration from native vasculature onto synthetic substrates, endothelial cells must transition from a migratory, proliferative phenotype to a quiescent monolayer that displays anti-thrombotic phenotypes for sustained thromboresistance.^27, 28^ As the endothelial cells migrate onto subsequent substrates, their initial attachment is established through integrins first binding to extracellular matrix ligands, followed by integrin activation and clustering at the cell membrane.^29–31^ Integrin clustering promotes intracellular signaling and focal adhesion assembly. ^30, 31^ Focal adhesion complexes serve as mechanosensitive signaling hubs that regulate downstream cellular functions, including migration, proliferation, and phenotypic state.^30, 31^ Biochemical cues like ligand identity and concentrations in addition to biophysical cues presented by the underlying material modulate integrin clustering and focal adhesion maturation, thereby directing cell behavior.^32, 33^ One of the critical ligands of interest is the GFOGER of collagen, a primary component of the basement membrane, which engages with integrins α1β1 and α2β1.^34, 35^ This interaction facilitates adhesion and migration across vascular constructs, making these integrins of particular relevance for biomaterial design.^36–39^ Although integrin engagement and focal adhesion formation in collagen-based endothelial cell behavior has been studied, the temporal regulation of integrin expression and focal adhesion formation during endothelialization on blood-contacting devices remains poorly understood.^12, 20, 40^ Previous work has largely characterized static endpoints rather than mapping how integrin expression profiles and associated focal adhesion structures develop over time in response to stiffness.^11, 12, 20, 40^ During transanastamotic endothelialization, cells must migrate from the native vessel onto synthetic graft materials, encountering a dramatic shift in mechanical environment as they transition from native vasculature to comparatively stiff substrates.^19, 41^ This change in stiffness directly modulates integrin expression, clustering, and downstream signaling, ultimately influencing focal adhesion maturation, cytoskeletal organization, and hemostatic phenotypes.^11, 29, 42, 43^ As a result, substrate stiffness governs the balance between stable adhesion and effective migration, both of which are essential for successful endothelialization.^44^ Understanding how endothelial cells dynamically regulate integrin expression and adhesions in response to these mechanical disparities is therefore critical, as mismatches in stiffness can impair endothelial coverage, delay healing, and contribute to thrombosis or graft failure.

Additionally, most *in vitro* investigations of endothelial cell behavior rely on preliminary expansion on treated tissue-culture polystyrene (TCPS) platforms with subsequent culture on experimental substrates.^45^ While this approach is widely adopted for ease of cell expansion, it introduces a mechanical conditioning step that can interfere with cell mechanical memory and change native cell phenotypes.^46, 47^ TCPS substrates possess moduli in the range of Gigapascals (∼ 1 GPa), which is significantly greater than any native arterial tissue stiffness.^48^ Because endothelial cell behavior is strongly correlated with substrate stiffness, culture on TCPS can impose bias on subsequent downstream pathways.^49, 50^ Consequently, *in vitro* interpretations of successful transanastamotic migration may not translate to *in vivo* device performance due to differences in cell substrate expansion and culture conditions. These considerations highlight the importance of investigating expansion and culture condition’s effect on endothelial cell behavior to better understand translation to transanastamotic migration conditions.

In this study, we explore the effect of substrate stiffness on integrin expression and the corollary effect on endothelial cell phenotype. Time-dependent integrin expression of α1 and α2 subunits was investigated for human coronary artery endothelial cells (HCAECs) seeded and reseeded on either TCPS or bioactive hydrogels of different substrate stiffnesses that match the ranges of native vessels, decellularized matrices, or synthetic conduits. These differences in integrin expression profiles were then correlated to focal adhesion formation, proliferation and quiescence, migratory behavior, and hemostatic phenotypes to impart relevance to integrin-mediated transanastamotic migration. Collectively, these studies will provide a mechanistic framework for understanding time-dependent changes in cell behavior based on culture conditions towards designing synthetic conduits with appropriate endothelial cell phenotypes.

## Methods

### Materials

Reagents were purchased from Sigma-Aldrich and used without further purification unless otherwise noted.

### Cell culture

Three male and three female donor HCAECs were purchased from Promocell. Donors were age-matched, ranging from 50 to 65 years old, to minimize the confounding effect of age on cell behavior. Each donor set was expanded using Endothelial Cell Growth Media (Promocell) and 20% fetal bovine serum (FBS). FBS was charcoal stripped to remove lipophilic molecules, including hormones, that can interfere with experimental studies. Cells were expanded in media with phenol red because of the increased proliferation rate; however, the media was replaced with phenol red-free media 24 hours before each study to limit estrogenic effects from the phenol red. All cultures were grown in tissue culture polystyrene flasks incubated at 37°C with 5% CO_2_. Media was exchanged every 2-3 days.

### Synthesis of Polyethylene Glycol Diacrylate

Polyethylene glycol diacrylate (PEGDA) was synthesized using the protocol from Browning et al. with minor alterations.^51^ Briefly, a solution of PEG (molecular weight 3.4 kDa, 10 kDa, or 20 kDa; 1 equivalent) was prepared in anhydrous dichloromethane under nitrogen. Triethylamine (2 molar equivalents) was added dropwise to the PEG solution, followed by dropwise addition of acryloyl chloride (4 molar equivalents). The solution was mixed for 24 hours. The reaction solution was then neutralized with potassium bicarbonate (8 mole equivalents) and dried with sodium sulfate. The polymer was precipitated in cold ether, vacuum filtered, and then dried under vacuum. Synthesis of PEGDA was confirmed using proton nuclear magnetic resonance (^1^H-NMR) spectroscopy. Proton NMR spectra were recorded on a Mercury 300 MHz spectrometer using a TMS/solvent signal as an internal reference. ^1^H-NMR (CDCl_3_): 3.6 ppm (m, -OC*H_2_*C*H_2_-*), 4.3 ppm (t, -C*H_2_*OCO-) 6.1 ppm (dd, -C*H*=CH_2_), 5.8 and 6.4 ppm (dd, -CH=C*H_2_*). Polymers with percentage conversions of hydroxyl to acrylate end groups over 85% were used in this work.

### Bioactive Hydrogel Fabrication

Bioactivity was imparted to PEGDA (3.4, 10, 20 kDa) hydrogels through incorporation of either Scl2-2 with an acrylamide-isocyanate linker (functionalization of ∼10% of available lysines).^51^ Similarly, bioactive thermoresponsive PEGDA-NIPAAm hydrogels (3.4, 10, 20 kDa PEGDA) with w/v ratio of NIPAm to that of PEGDA was 4%:16% PEGDA-NIPAAm (3.4 or 10 kDa PEGDA) and 3%:6% PEGDA-NIPAAm (20 kDa PEGDA). Bioactive hydrogels were prepared by dissolving PEGDA and functionalized Scl2-2 (8 mg/mL) in 50 mM acetic acid. A photoinitiator solution of Irgacure 2959 (10 wt/vol% solution in 70 vol% ethanol) was added to precursor solutions at a final concentration of 0.1 wt/vol%. Hydrogel sheets were fabricated by pipetting precursor solutions between 0.75 mm spaced glass plates and curing on a UV transilluminator (UVP, 25 watt, 365 nm) for 6 minutes on both sides.

### Stiffness Characterization

Hydrogel discs (D = 8 mm) samples were punched from hydrogel sheets and swollen to equilibrium. Samples were subjected to mechanical testing using a dynamic mechanical analyzer (RSAIII, TA Instruments) equipped with a parallel-plate compression clamp. Testing was performed under unconstrained compression at 37 C. Dynamic strain sweeps were used to determine the linear viscoelastic range for each hydrogel formulation. Compressive modulus was determined from a frequency sweep at 1% strain. The compressive storage modulus was taken at 0.5 Hz in the linear region (n = 12 specimens).

### Flow Cytometry

Differences in integrin expression between the male and female HCAECs was determined using antibody staining for flow cytometry. HCAECs were cultured in tissue culture polystyrene flasks to 90% confluency. Cells were then either reseeded on TCPS substrates or cultured on PEG-NIPAam-Scl2 hydrogels of different substrate stiffnesses for 24 hours, 3 days, or 1 week. Cells were lifted from TCPS substrates with accutase or retrieved from PEG-PNIPAAm-Scl2 substrates by placing substrates between 4 C and 37 C every 15 minutes for 2 hour. Lifted cells were then spun down at 220 rcf followed by resuspension in flow buffer (2% FBS in phosphate buffered saline (PBS)) at a concentration of 10^5^ cells/ml. Each sample was stained with fixable viability dye 450 (BD Biosciences) on ice for 20 minutes to selectively analyze viable cells. After washing samples with 3 mL of flow buffer, cells were centrifuged and resuspended in flow buffer. Samples were titrated with their respective phycoerythrin-conjugated integrin antibodies (anti-α1 (BioLegend anti-human CD49a), anti-α2 (BioLegend anti-human CD49b)) in 0.5 ug increments on ice to identify signal saturation. Each sample was then washed with 3 mL of flow buffer and spun down. Cells were resuspended in 400 uL 4% paraformaldehyde for fixation for 15 minutes. Blank samples of cells without antibodies underwent similar protocols. Samples were then rinsed with 3 mL flow buffer, spun down, and resuspended in 1 mL flow buffer. An Attune Nxt Acoustic Focusing flow cytometer was used to evaluate surface expression of integrins. Relative fluorescence of each integrin was assessed with each donor. For each integrin tested, samples were run in triplicate (6 biological donors, n = 3 technical replicates per donor). Fluorescence intensity was analyzed using FlowJo software. The combined average of all biological and experimental replicates is represented for each donor.

### Endothelial Cell Proliferation

HCAECs were seeded on bioactive hydrogels at 15,000 cells/cm^2^ on hydrogel discs (n = 3 for donor) and allowed to attach for 24 hours, 3 days, or 1 week. Cells were then washed twice with PBS to remove non-adherent cells. Specimens were then fixed with 3.7% glutaraldehyde and stained with SYBR green (DNA/nucleus) for 10 minutes then specimens followed by two washes with PBS. Specimens were imaged (3 images per specimen and *n* = 3 specimens per donor for a total of 9 images per donor) using a fluorescence microscope (Nikon Eclipse TS100) to quantify cell adhesion and proliferation. The average adhesion and spreading of each donor were reported. Briefly, cell adhesion was assessed by manually counting the number of SYBR Green-stained nuclei for each image. The combined average of all experimental replicates is represented for each donor (n = 3 donors per sex).

### Immunofluorescence Staining for Integrins and Focal Adhesions

Cells seeded for 1 week on bioactive hydrogels were fixed to gel specimens with 4% paraformaldehyde then permeabilized with 0.1% Triton X-100. Permeabilized cells then underwent blocking with 1% BSA in PBS for 30 minutes. Cells were then incubated with either anti-vinculin primary antibody, anti-α1 primary antibody, or anti-α2 primary antibody for 24 h, followed by three washes with wash buffer. A FITC-conjugated goat anti-mouse secondary antibody (e.g., Millipore, Cat. No. AP124F) was then incubated for 60 min for anti-vinculin samples. An AlexaFluor88-conjugated goat anti-rabbit secondary antibody (e.g., Millipore, Cat. No. AP124F) was then incubated for 60 min for anti-integrin samples. TRITC-conjugated phalloidin was incubated concurrently with the secondary antibody. Nuclei were counterstained with DAPI for 5 min. Fluorescence images were acquired using a confocal microscope (Leica SP8 Confocal Microscopy).

### Cell Migration

Endothelial cell migration was assessed following seeding on hydrogels stamped with a custom mold fabricated from Sylgard 182 polydimethylsiloxane (PDMS) to create a cell-free gap (0.5 mm wide). After achieving confluence, the mold was removed to create a defined linear gap to initiate migration. Brightfield images were collected immediately following mold removal (t = 0 h) and at subsequent 24 h time points to monitor gap closure. Migration was quantified by measuring the gap area percentage over time using ImageJ.

### Time to Confluence

Time to confluence was evaluated on bioactive hydrogels. Specimens were seeded at 20,000 cells/cm². Brightfield images were captured every 24 h until confluence was reached, and confluence was defined as the first timepoint at which ≥ 90% of the hydrogel surface area was covered by cells (3 images per specimen and *n* = 3 specimens per donor for a total of 9 images per donor). Percent confluence was scored using a Matlab code designed to evaluate confluency.

### Ki-67 Detection Assay

Cell quiescence and proliferation was quantified using the Lumit® Ki-67 Immunoassay (Promega) according to the manufacturer’s protocol. Briefly, cells were lysed by addition of 20 µL of 5X Lumit® Lysis Buffer II per well and incubated with shaking at 800 rpm for 40 minutes at room temperature. A 2X antibody mixture containing Anti-hKi-67 mAb-SmBiT and Anti-hKi-67 mAb-LgBiT was added at 100 µL per well and incubated for 90 minutes at room temperature. Lumit® Detection Reagent C was then added at 50 µL per well, and luminescence was recorded after a 3–5 minute incubation (n = 3 specimens per substrate condition).

### Nitric Oxide Detection

Total nitric oxide (NO) production was assessed by measuring nitrate/nitrite concentrations of cell media following 1 week of culture using the Nitric Oxide Assay Kit (MAK454, Sigma-Aldrich). Media samples underwent deproteination by mixing 150 µL of sample with ZnSO and NaOH, followed by centrifugation and supernatant retention. A nitrite standard curve was generated by diluting a 100 µM nitrite working standard to concentrations of 0, 30, 60, and 100 µM, with all samples and standards run in duplicate. Working reagent was added to each sample and standard then incubated at 60 °C for 10 minutes. The optical density was measured at 540 nm using a spectrophotometric plate reader with each sample run in triplicate.

### Statistical Analysis

The data for all other measurements are displayed as mean ± standard deviation. Statistical comparisons were made using an analysis of variation (ANOVA) comparison utilizing Tukey’s post-hoc analysis for parametric data. Computations were performed using GraphPad Prism version 10 at the significance levels of p < 0.05.

## Results

### Effects of Hydrogel Stiffness on Integrin-Mediated Endothelialization Processes over Time

To first determine how substrate stiffness regulates α1β1 and α2β1 integrin expression and focal adhesion maturation, integrin subunit expression in HCECs initially expanded on TCPS and subsequently reseeded onto hydrogels of varying stiffness was quantified by flow cytometry. Across all conditions and time points, α1 expression remained consistently lower than α2 expression (**Figure 1a,b**), consistent with previous reports demonstrating lower α1 expression relative to α2 in endothelial cells cultured on TCPS-expanded conditions.^12^ Importantly, α1 subunit expression did not exhibit significant stiffness-dependent or temporal regulation, suggesting α1β1 engagement is relatively insensitive to substrate stiffness or time within the tested range. In contrast, α2 subunit expression demonstrated both substrate- and time-dependent effects, indicating α2β1 may represent the dominant stiffness-responsive collagen-binding integrin in this system. At 24 h timepoints, hydrogels with 300 kPa stiffnesses showed the highest level of α2 expression, followed by 100 kPa and 20 kPa hydrogels, respectively. Similar trends were maintained through 3 day and 1-week timepoints, with statistically significant increases in α2 subunit expression between softest (20 kPa) and stiffest (300 kPa) substrate stiffness. In addition, soft and intermediate substrates stiffnesses do not significantly increase in integrin expression over time (**Figure S1**). However, stiff substrates substantially increase in α2 subunit expression between 24h and 3 days. Integrin expression trends from flow cytometry were validated by confocal imaging, which revealed similar increases in α2-associated signal intensity between soft and stiff substrates at 1-week timepoints (**Figure S4**). Consistent with the flow cytometry results, α1 immunofluorescence signal was near the detection limit and could not be clearly resolved by confocal microscopy across all substrate stiffnesses, further supporting the relatively low abundance of α1 in HCECs. Immunostaining for focal adhesion protein vinculin and actin were used to confirm focal adhesion formation. Enhanced vinculin intensity and actin cytoskeletal area were observed on stiff substrates as compared to soft substrates, indicative of increased focal adhesion maturation (**Figure 1c)**. Visualization of actin fibers also revealed greater cytoskeletal organization with increasing substrate stiffness. Together, these results demonstrate that increasing substrate stiffness preferentially enhances α2β1-associated mechanotransduction and focal adhesion maturation, suggesting α2β1-mediated interactions are major regulators of endothelial cell responses to stiff biomaterial environments.

**Figure 1:**
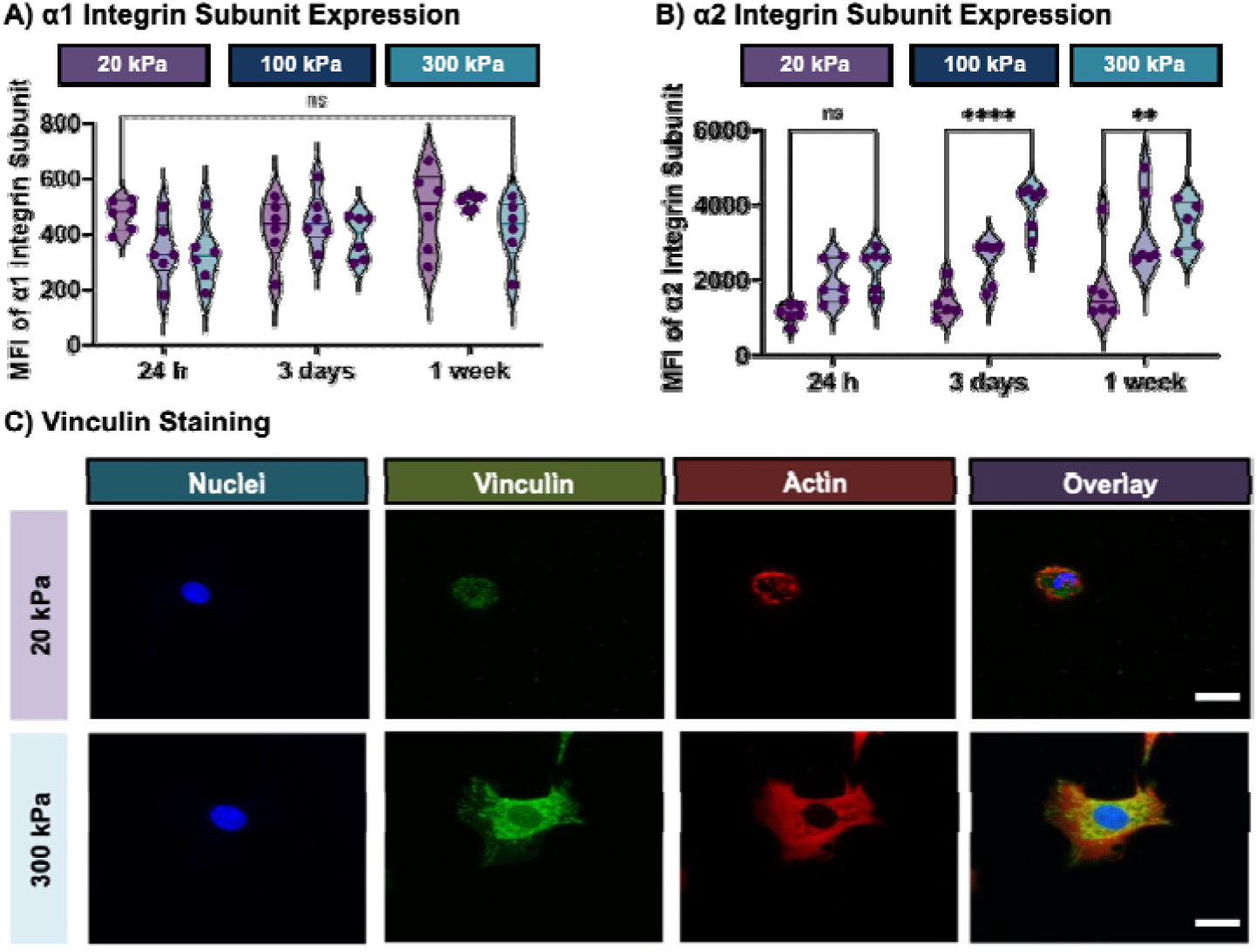
Effect of substrate stiffness on endothelial cell integrin expression and focal adhesion formation on PEG-Scl2 hydrogels (20, 100, or 300 kPa). A, B) Time-dependent changes in α1 and α2 integrin expression after 24 h, 3 d, and 1 week of culture on PEG-Scl2 hydrogels quantified by flow cytometry. Data represents six biological donors, with each data point corresponding to the average of three technical replicates per donor. C) Representative immunofluorescence images of the focal adhesion protein vinculin after 1 week of culture on PEG-Scl2 hydrogels (20 or 300 kPa) (n = 3 specimens per donor, 8 images per specimen) (scale bar = 10 µm). Vinculin was stained using an anti-vinculin antibod with FITC to assess focal adhesion formation. Data are presented as mean ± standard deviation across all donors (** = p < 0.01; **** = p < 0.0001; ns = not significant).

Following identification of stiffness-dependent α2β1 integrin upregulation at all timepoints and focal adhesion maturation on stiff substrates, subsequent studies investigated whether these changes corresponded with alterations in behaviors associated with endothelialization, including density, migration, confluence, and quiescence. Initial cell attachment and subsequent proliferation increased with substrate stiffness, with significantly higher cell densities observed on stiffer substrates at later time points (**Figure 2a)**. Similarly, endothelial cell migration increased with substrate stiffness, as cells cultured on stiffer hydrogels exhibited greater motility than those cultured on softer hydrogels (**Figure 2b)**. In fact, cells on the softest substrate exhibited remain stagnant with no minimal changes in migration over time. Time-to-confluence analysis further supported these findings, as endothelial cells cultured on stiffer substrates reached confluence more rapidly and at greater extents (**Figure 2c)**. Cells cultured on stiff substrates exhibited the most rapid increase in confluence, reaching approximately 90% by day 6, whereas cells on 100 kPa substrates increased more gradually and plateaued at approximately 55% by day 4. In contrast, cells cultured on 20 kPa substrates remained sparsely distributed throughout the study, with confluence remaining below 15% at all time points. To determine whether stiffness-dependent proliferation and migration persisted long-term or ultimately transitioned toward stabilized endothelial phenotypes, Ki67 was used to assess quiescent phenotypes. Cells cultured on stiff substrates demonstrated higher concentrations of Ki67 at 24 h timepoints, indicating enhanced proliferative phenotypes in comparison to intermediate and soft substrates at the same timepoint (**Figure 2d)**. However, assessment of quiescent states via the absence of Ki-67 revealed a temporal shift toward quiescence, with a decrease in Ki-67–positive cells over time across all substrate conditions. After one week in culture, cells on each substrate exhibited predominantly quiescent phenotypes, indicating that despite initial increases in proliferation and migration on stiff substrates, endothelial cells ultimately transition to a stable, non-proliferative state regardless of substrate stiffness. These results suggest that while substrate mechanics modulate cell growth and motility, endothelial quiescence remains relatively stable across the tested stiffness range after 1 week.

**Figure 2:**
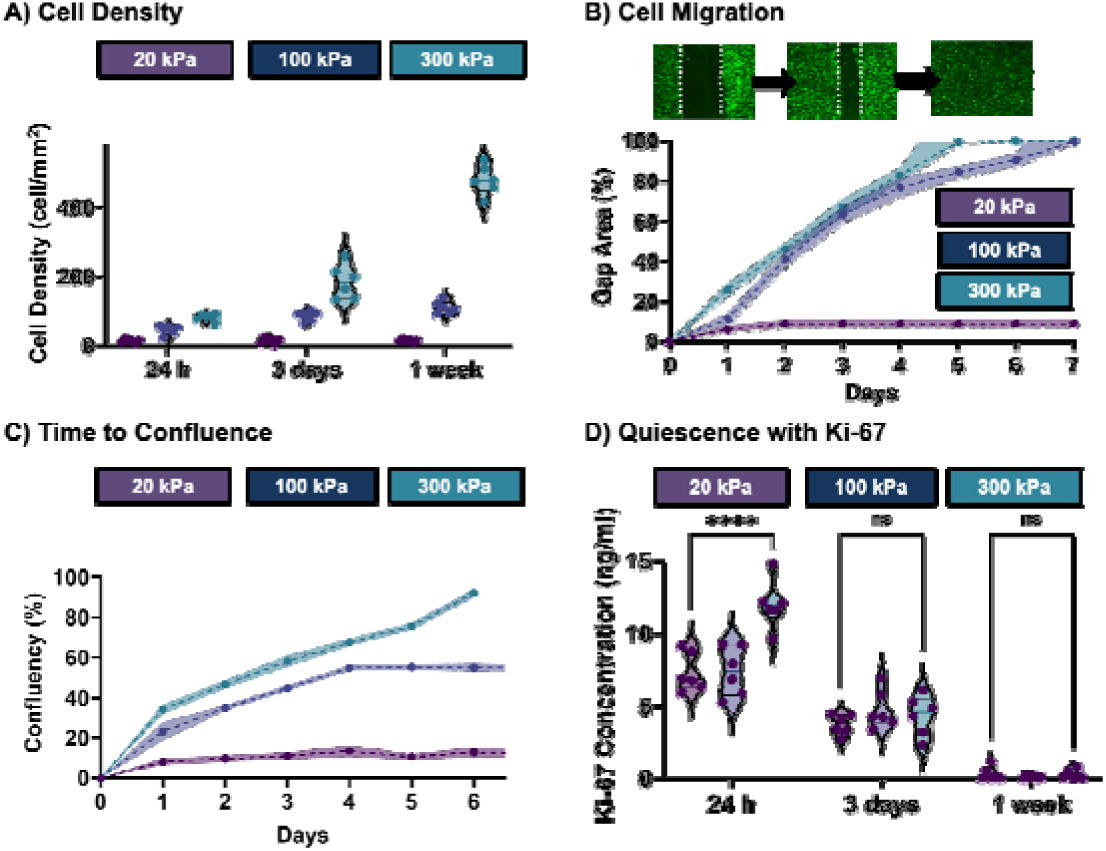
Effect of substrate stiffness on endothelial cell density, migration, confluence, and quiescenc on PEG-Scl2 hydrogels (20, 100, or 300 kPa). A) Endothelial cell density quantified over time on PEG-Scl2 hydrogels (6 donors, n = 3 specimens per donor, 3 images per specimen). B) Cell migration assessed as a function of substrate stiffness and culture duration. C) Time to confluence of endothelial cells cultured on PEG-Scl2 hydrogels of varying stiffness. D) Ki-67 expression used to evaluate proliferative and quiescent cell phenotypes across substrate stiffness conditions (6 donors, n = 3 specimens per donor). Data represent six biological donors, with each data point corresponding to the average of three technical replicates per donor. Data are presented as mean ± standard deviation across all donors (*** = p < 0.001; ns = not significant).

### Effects of Expansion Conditions on Integrin-Mediated Endothelialization Processes

To evaluate whether expansion condition creates a persistent mechanical memory effect that alters future integrin responses, integrin subunit expression was quantified in HCECs initially expanded on soft hydrogels and subsequently reseeded onto stiffer hydrogel environments (300 kPa). Similar to TCPS-expanded conditions, α1 expression remained consistently lower than α2 expression for the gel expansion condition at all time points. In addition, α1 expression demonstrated minimal dependence on expansion condition or culture duration, showing that α1β1 engagement is relatively insensitive to expansion condition (**Figure 3a).** In contrast, α2 integrin expression varied over time as a function of cell expansion conditions (**Figure 3b**). At 24 h, hydrogel-expanded cells exhibited significantly lower α2 integrin expression than TCPS-expanded cells. By 3 days, α2 integrin expression increased in both groups, with TCPS-expanded cells displaying significantly greater expression than hydrogel-expanded cells. At 1 week, α2 integrin expression remained elevated in both expansion conditions with no significant differences were observed between TCPS- and hydrogel-expanded cells. Overall, the differences in α2 integrin expression associated with expansion conditions were most pronounced during the early culture period and diminished by 1 week. In agreement with the trends seen with flow cytometry, confocal imaging demonstrated similar levels of α2-associated intensity at the same substrate stiffness after 1 week in comparison to cells initially cultured on TCPS (**Figure S7**). In addition, immunostaining for vinculin demonstrated similar levels of intensity regardless of initial culture substrate (**Figure 3c)**. Actin staining similarly revealed no significant differences in cell spreading or cytoskeletal organization across conditions.

**Figure 3:**
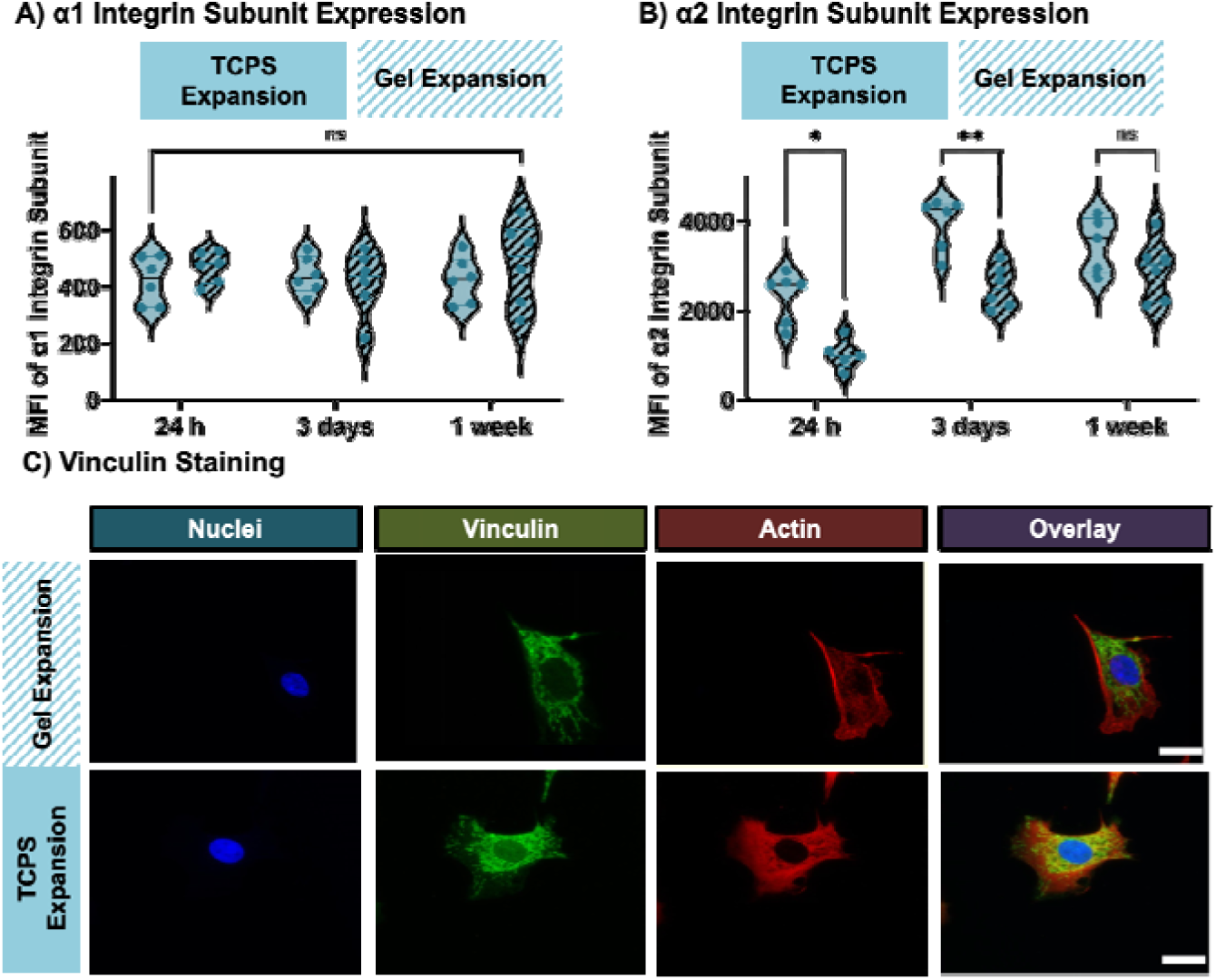
Effect of substrate expansion condition (TCPS or soft hydrogel) on endothelial cell integrin expression and focal adhesion formation on stiff PEG-Scl2 hydrogels (300 kPa). A, B) Time-dependent changes in α1 and α2 integrin expression after 24 h, 3 d, and 1 week of culture on PEG-Scl2 hydrogel quantified by flow cytometry. Data represents six biological donors, with each data point corresponding to the average of three technical replicates per donor. C) Representative immunofluorescence images of the focal adhesion protein vinculin after 1 week of culture on PEG-Scl2 hydrogels (n = 3 specimens per donor, 8 images per specimen). Vinculin was stained using an anti-vinculin antibody with FITC to assess focal adhesion formation. Data are presented as mean ± standard deviation across all donors (* = p < 0.05; ** = p < 0.01; ns = not significant).

Cell density and migration of HCECs expanded on soft hydrogels matching native vessel stiffnesses and then transferred to stiff (300 kPa) substrates were examined over time to determine how mechanical priming on substrates matching native vessel stiffnesses influence integrin-mediated endothelial behavior. Adhesion and proliferation increased with culture duration for both expansion conditions with no statistically significant differences in proliferation **(Figure 4a)**. Cells expanded on hydrogel substrates also displayed migration profiles similar to those of cells cultured on TCPS and reseeded onto substrates of the same stiffness at the same timepoint, with only a slight reduction in migration following reseeding onto hydrogels **(Figure 4b)**. Analysis of time to confluence further aligned with these observations, as cells initially seeded on TCPS reached confluence at similar rates to cells expanded on hydrogels (**Figure 4c)**. Evaluation of quiescence demonstrated a time-dependent decline in proliferative phenotypes for both substrate conditions (**Figure 4d)**. By one week, cultures from both expansion conditions were largely Ki-67–negative, indicating the conferral of quiescent phenotypes regardless of expansion condition.

**Figure 4:**
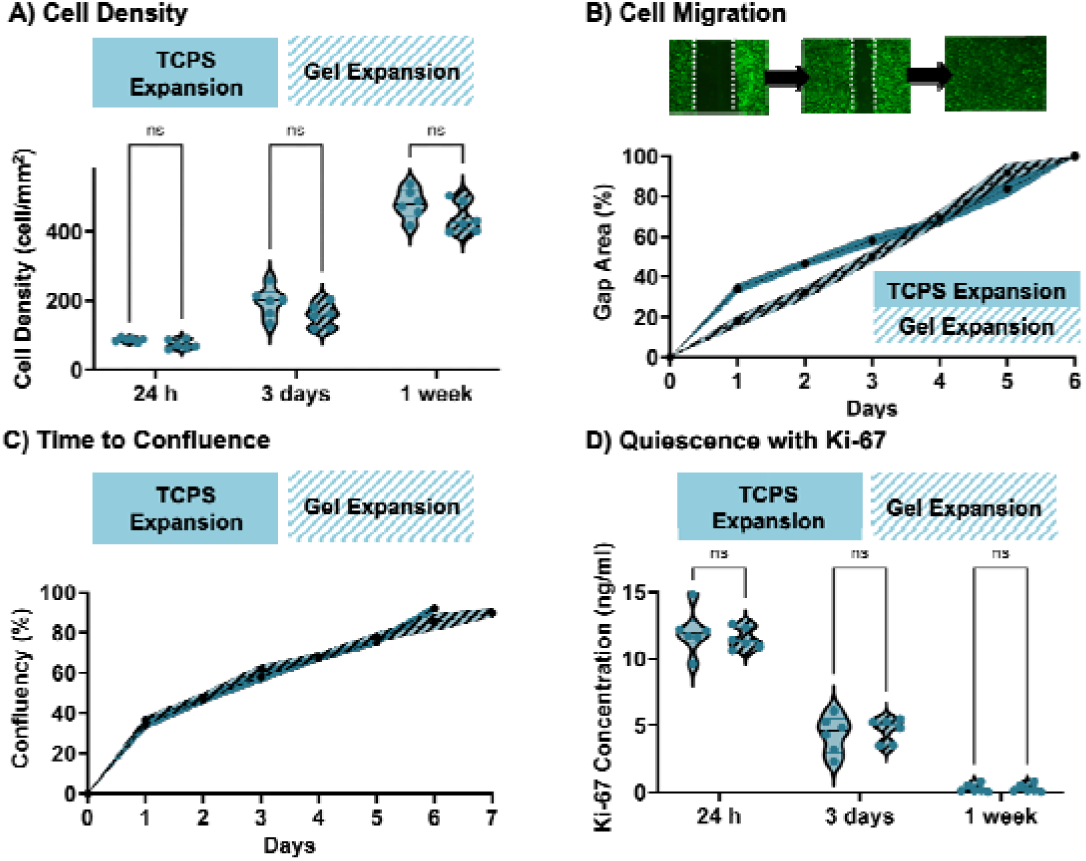
Effect of substrate expansion condition (TCPS or soft hydrogel) on endothelial cell density, migration, confluence, and quiescence on stiff PEG-Scl2 hydrogels (300 kPa). A) Endothelial cell densit quantified over time on PEG-Scl2 hydrogels (6 donors, n = 3 specimens per donor, 3 images per specimen). B) Cell migration assessed as a function of expansion condition and culture duration C) Time to confluence of endothelial cells cultured on PEG-Scl2 hydrogels. D) Ki-67 expression used to evaluate proliferative and quiescent cell phenotypes across substrate expansion conditions (6 donors, n = 3 specimens per donor). Data represent six biological donors, with each data point corresponding to the average of three technical replicates per donor. Data for all comparisons represents the combined average of each biological and experimental replicate per donor (ns = not significant).

## Discussion

Endothelialization is vital to ensure long-term thromboresistance of blood-contacting devices.^3, 11^ Towards this objective, successful deployment of biomaterials for blood-contacting devices is contingent on the conferral of a confluent endothelial cell layer to promote anti-thrombotic phenotypes.^52^ Previous studies have investigated integrin-mediated endothelial cell behavior on a variety of extracellular matrix (ECM) proteins and biomimetic substrates towards conferral of this thromboresistant monolayer; however, they have largely failed to isolate the individual and combined effects of substrate stiffness, time, and expansion condition on integrin expression and subsequent integrin-mediated endothelial cell function.^20, 51^ Therefore, PEG-Scl2 hydrogels were used as a platform to selectively isolate the relative contributions of α1β1 and α2β1 integrin-mediated phenotypes using different substrate stiffnesses, as these collagen-binding integrins play a central role in mediating endothelial adhesion, mechanotransduction, and downstream regulation of cell phenotype.^11^

Because these processes are largely mediated through integrin signaling, clustering, and focal adhesion assembly, characterization of integrin expression and focal adhesion complex formation is an important first step in understanding endothelial cell responses to biomaterial environments.^53^ Although α1 subunit expression remained relatively unchanged across all stiffnesses and timepoints, α2 subunit expression exhibited clear stiffness- and time-dependent trends, suggesting that α2β1 integrins likely plays a more dominant role in endothelial mechanosensing. Although α2β1 expression generally trended upward with increasing substrate stiffness, statistically significant differences were observed between the softest and stiffest substrates at 3 days and 1 week. This trend suggests that endothelial cells dynamically adjust integrin-mediated adhesion in response to increasing matrix rigidity. Consistent with this concept, Yeh et al. demonstrated that β1 integrin expression is dynamically regulated by matrix stiffness, with increasing β1 integrin accumulation on stiff substrates over time, supporting the ability of endothelial cells to adapt their integrin-ligand interactions in response to mechanical cues.^43^ Minimal temporal changes were observed on the softest hydrogels, whereas α2β1 expression increased substantially on the intermediate and stiff substrates, with the most pronounced change occurring between 24 h and 3 days for stiff substrates. Integrin α2β1 has been associated with enhanced cytoskeletal organization, force transmission, and focal adhesion assembly, all of which are critical for endothelial spreading and stabilization on synthetic materials.^54^ This was further validated with confocal imaging where vinculin staining revealed enhanced focal adhesion intensity on stiff hydrogels relative to soft hydrogels, indicative of increased focal adhesion maturation. This increase in focal adhesion formation after 1 week in culture was temporally associated with elevated α2 expression at later timepoints, suggesting that α2β1 upregulation may contribute to the reinforcement of focal adhesion structures. Substrate rigidity is known to regulate focal adhesion formation through force-dependent stabilization of integrin–cytoskeletal linkages, enabling recruitment of proteins such as vinculin, talin, and paxillin into mature adhesion complexes.^55^ The increased vinculin intensity observed therefore suggests that stiffer substrates promote stronger and more stable cell–matrix interactions. Because integrin expression and focal adhesion maturation is closely linked to endothelial migration, spreading, and proliferation, these findings collectively indicate that stiff substrates may promote a more adhesive and mechanically reinforced endothelial phenotype.^11, 56, 57^

These findings were further validated through assessments of integrin-mediated adhesion, proliferation, migration, and confluence. HCECs cultured on stiffer hydrogels exhibited enhanced proliferation, increased migration, and more rapid progression toward confluence compared to cells on softer substrates, with these differences becoming increasingly pronounced over time. This trend is consistent with previous studies demonstrating that increasing substrate stiffness promotes endothelial cell activation through enhanced cytoskeletal tension, focal adhesion maturation, and mechanotransductive signaling, ultimately driving increased proliferation, migration, and tissue remodeling responses.^58, 59^ In contrast, cells cultured on soft hydrogels showed minimal temporal changes in proliferation, migration, or confluence, suggesting that their functional phenotype remained relatively static throughout the culture period. Notably, this limited functional response paralleled the absence of significant changes in either α1β1 or α2β1 integrin expression on soft substrates, whereas stiff substrates induced a time-dependent upregulation of α2β1 expression accompanied by increased vinculin staining. This finding is consistent with previous studies demonstrating that α2β1-mediated adhesion is a major contributor to endothelial cell attachment to PEG-collagen and PEG-Scl2 substrates, with inhibition of α2 integrins resulting in more pronounced reductions in cell adhesion compared with α1 integrin blockade. Specifically, blocking α2 integrins produced a greater decrease in endothelial cell attachment than α1 integrin inhibition, highlighting the dominant role of α2β1 signaling in mediating cell–matrix interactions.^12^ Because α2β1 integrins are key mediators of endothelial cell adhesion and mechanotransductive signaling, enhanced α2β1 engagement on stiff hydrogels likely promoted focal adhesion assembly and cytoskeletal organization, strengthening cell–matrix interactions and facilitating the force transmission required for migration and proliferation. ^54, 60^ The associated increases in vinculin intensity further indicates greater focal adhesion maturation and stabilization, supporting the formation of mechanically reinforced adhesive structures capable of sustaining enhanced cell proliferation and migration on stiff substrates. Together, these findings suggest that stiffness-mediated increases in α2β1 signaling and focal adhesion maturation may represent key mechanistic drivers underlying the enhanced proliferative and migratory endothelial phenotype observed on stiff substrates. Importantly, Ki-67 analysis further demonstrated that while endothelial cells cultured on stiff substrates exhibited increased proliferative activity during early culture, Ki-67 expression decreased across all substrate conditions over time, with cells reaching a predominantly quiescent phenotype by 1 week. This temporal shift in Ki-67 expression is consistent with previous studies demonstrating that Ki-67 levels dynamically fluctuate with cell-cycle progression and decline as cells enter quiescence.^61^ Thus, the early increase in Ki-67 observed on stiff substrates likely reflects enhanced proliferative activity during initial endothelialization, whereas the subsequent reduction across all conditions suggests stabilization of quiescent phenotypes. Collectively, these findings demonstrate that substrate stiffness regulates endothelialization dynamics by enhancing α2β1 integrin-mediated mechanotransduction and focal adhesion maturation on stiff substrates, thereby accelerating endothelial proliferation and migration while still allowing progression toward a stable, quiescent endothelial phenotype over time.

Although TCPS is widely used for endothelial cell expansion, it does not recapitulate the mechanical environment encountered during transanastomotic migration, where endothelial cells transition from the native vessel wall onto the surface of implanted blood-contacting devices.^25, 47^ Therefore, evaluating endothelial responses following initial expansion on substrates that mimic native vessel stiffness and subsequent transition to substrates representative blood-contacting devices such as hydrogel-coated implants provides a more physiologically relevant framework for understanding how mechanical history influences integrin-mediated endothelial behavior during migration. Given that endothelial cells experience dynamic changes in matrix stiffness throughout implantation, determining how prior substrate exposure affects integrin expression and subsequent adhesion, proliferation, and migration responses is critical for designing materials that support effective endothelialization of blood-contacting devices.^20, 62^ Despite the distinct mechanical environments experienced during initial expansion, prior culture conditions did not produce sustained differences in integrin-mediated endothelial responses following transition onto hydrogel substrates. Across all timepoints, no statistically significant differences in α1 integrin expression were observed between TCPS-expanded and hydrogel-expanded cells. Interestingly, TCPS-expanded cells exhibited increased α2β1 integrin expression at early timepoints (24 h and 3 days) compared to hydrogel-expanded cells, suggesting that expansion on a stiffer substrate may temporarily prime endothelial adhesion signaling through increased integrin engagement. However, this difference was no longer apparent after 1 week, indicating that α2β1 expression was progressively regulated by the mechanical cues of the current hydrogel environment rather than maintained through prior substrate exposure. This observation aligns with previous findings demonstrating that endothelial cells expanded on mechanically distinct substrates undergo early stiffness-dependent changes in adhesion signaling and functional state, reflecting an initial adaptive response to changes in the extracellular mechanical environment.^47^ This temporal adaptation is consistent with the dynamic nature of integrin-mediated mechanotransduction, where endothelial cells continuously remodel their adhesion profiles in response to changes in extracellular matrix properties.^63, 64^ Importantly, the elevation in α2β1 expression observed during the first 3 days did not result in downstream differences in focal adhesion maturation, as vinculin staining revealed comparable focal adhesion profiles between expansion conditions after 1 week. Furthermore, the absence of differences in adhesion, proliferation, migration, and progression toward quiescence at all assessed timepoints suggests that the early α2β1 upregulation in TCPS-expanded cells was insufficient to induce lasting functional changes. Together, these findings suggest that while prior exposure to a stiff TCPS environment can transiently influence α2β1 integrin expression during the early stages of substrate transition, endothelial cells rapidly adapt to their new mechanical surroundings and converge toward similar adhesive and functional phenotypes over time. This rapid mechanoadaptation indicates that the current extracellular environment encountered during migration may be a stronger determinant of endothelial behavior than expansion history, supporting the importance of designing blood-contacting materials that provide appropriate mechanical and biochemical cues during endothelialization.

### Study Limitations and Future Work

Although this study provides important insight into how substrate mechanics regulate endothelial integrin expression and downstream endothelial phenotypes relevant to transanastomotic migration, several limitations should be considered in the context of how these findings translate to vascular graft design. A key limitation of the current study is that endothelial responses were evaluated under static culture conditions and did not incorporate physiological shear forces. Since endothelial mechanotransduction is highly influenced by fluid shear stress, substrate stiffness-dependent changes in integrin expression, focal adhesion maturation, proliferation, and migration may differ under flow conditions.^63, 64^ Future studies using flow-based systems will be necessary to determine how mechanical cues from both the substrate and the vascular environment interact to regulate endothelialization. Furthermore, although this study focused on endothelial attachment, migration, and proliferation, successful conferral of thromboresistant phenotypes requires regulation of inflammatory responses and maintenance of a functional, antithrombotic endothelial phenotype. Future studies evaluating inflammatory markers and functional temporal measures of hemostatic regulation will be important to determine whether stiffness-dependent integrin signaling translates to improved vascular function. Incorporating more physiologically representative multicellular models, such as endothelial–smooth muscle cell and endothelial–platelet co-culture systems, will also be critical for evaluating how cell–cell interactions influence inflammatory and blood-contacting responses at engineered vascular interfaces. In addition, the hydrogel system used here was intentionally designed to isolate the effects of substrate stiffness and collagen-binding integrin engagement on endothelial behavior. By limiting the system to a single ligand concentration and defined stiffness conditions, we were able to directly connect increases in substrate stiffness with elevated α2β1 expression, enhanced vinculin-associated focal adhesion maturation, and subsequent increases in endothelial proliferation and migration. However, native vascular environments are substantially more complex as they contain dynamic mechanical gradients and multiple extracellular matrix ligands. As a result, endothelial mechanosensitive responses in vivo are likely regulated by the combined integration of these cues rather than stiffness alone. Future studies incorporating broader stiffness ranges and varied ligand densities will therefore be important to determine how these integrin-mediated responses evolve within more complex vascular environments. Similarly, the use of Scl2-functionalized hydrogels enabled more selective interrogation of α1β1- and α2β1-mediated interactions, allowing clearer mechanistic interpretation of how collagen-binding integrins contribute to stiffness-dependent endothelial phenotypes. This approach was particularly important for linking increased α2β1 expression with enhanced focal adhesion maturation and migratory behavior on stiff substrates. However, because native extracellular matrix and clinically used biomaterials contain numerous adhesive ligands capable of engaging multiple integrin populations simultaneously, endothelial responses in vivo may involve additional signaling pathways beyond those captured in the present study. Thus, although the hydrogel system strengthened mechanistic understanding of α2β1-associated mechanotransduction, it may not fully recapitulate the complexity of endothelial adhesion on other vascular graft materials in vivo. Finally, TCPS was used as a rigid control to mimic the substantial mechanical mismatch encountered during endothelial migration from compliant native vessels onto clinically relevant synthetic graft materials. While this comparison helped contextualize how endothelial cells adapt to rigid environments, TCPS does not allow selective interrogation of individual integrin interactions. Consequently, the observed endothelial responses on TCPS likely reflect combined adhesion signaling pathways rather than isolated α1β1- or α2β1-mediated effects. Together, these limitations highlight that the current findings should be interpreted as a mechanistic framework linking substrate stiffness, α2β1-associated mechanotransduction, focal adhesion maturation, and endothelialization-associated behaviors, rather than as a complete representation of the full in vivo vascular environment.

## Conclusion

In conclusion, this work demonstrates that substrate mechanics and time are a key regulator of endothelial integrin expression, focal adhesion maturation, and downstream behaviors relevant to transanastomotic migration. Using a mechanically defined system spanning soft, vessel-like hydrogels and stiff, graft-mimetic hydrogels or TCPS, endothelial cells were shown to undergo time-dependent shifts in adhesion and migration, including an increased reliance on α2β1-mediated adhesion and the development of more mature, vinculin-rich focal adhesions on stiffer substrates. These mechanosensitive changes were accompanied by enhanced proliferation and migration, followed by a transition toward a more quiescent endothelial phenotype over extended culture. Overall, these findings underscore the importance of mechanical history and substrate stiffness in shaping integrin-mediated endothelial behavior and provide mechanistic insight into how cells adapt during transanastomotic migration. This work advances our understanding of how endothelial cells integrate mechanical cues through integrin-mediated mechanisms to regulate adhesion, migration, and hemostatic function over time, key phenotypes underlying successful transanastomotic migration. These findings also provide mechanistic insight relevant to the design of blood-contacting devices, for which effective transanastomotic endothelialization is essential for long-term device integration and thromboresistance.

## Acknowledgments

This work was supported by the National Institutes of Health grant number R01 HL180615 and the National Science Foundation Graduate Research Fellowship Program under Grant (2023355666). Any opinions, findings, and conclusions or recommendations expressed in this material are those of the author(s) and do not necessarily reflect the views of the National Science Foundation.

