## Supplemental Information for "Mechanical History and Substrate Stiffness Shape Integrin-Mediated Endothelial Cell Behavior on Bioactive Hydrogels"

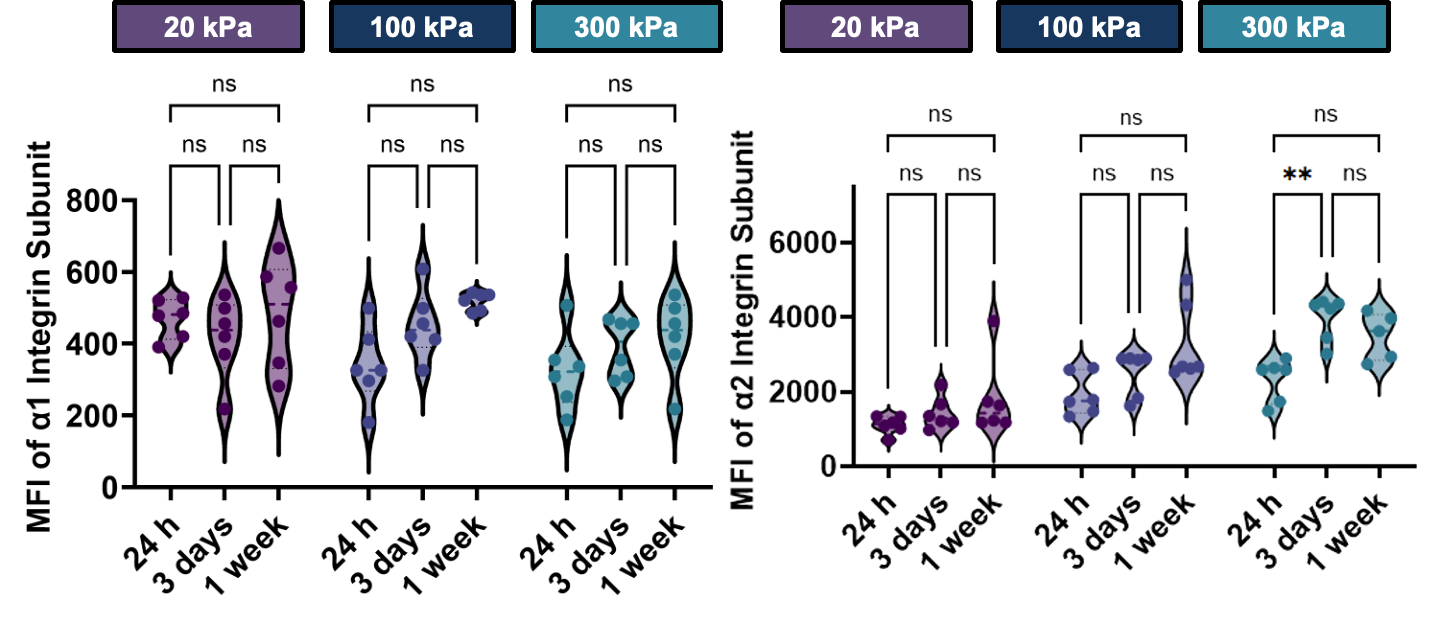

**Figure S1:** Temporal effects of substrate stiffness on endothelial α1 and α2 integrin expression. α1 and α2 integrin expression were quantified by flow cytometry after 24 h, 3 d, and 1 week of culture on PEG-Scl2 hydrogels with stiffnesses of 20, 100, or 300 kPa. Data represent six biological donors, with each data point corresponding to the average of three technical replicates per donor. Data are presented as mean ± standard deviation across all donors (** = p < 0.01; ns = not significant).

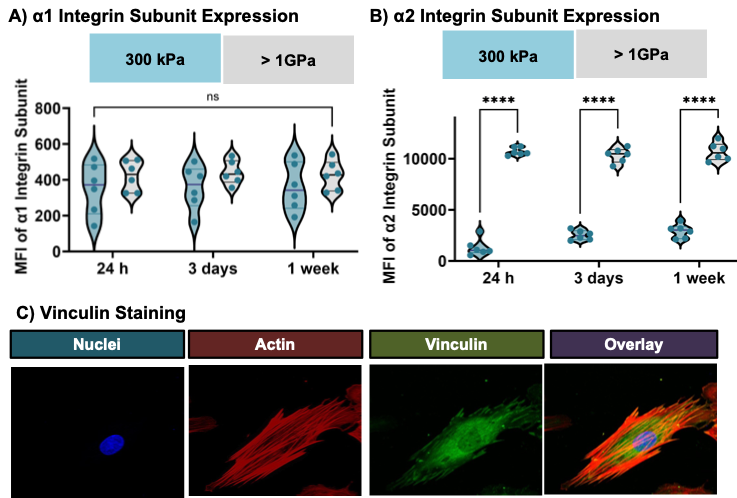

**Figure S2:** Effect of soft hydrogel expansion to stiff condition reseeding (1 GPa TCPS or 300 kPa hydrogel) on endothelial cell integrin expression and focal adhesion formation on PEG-Scl2. A, B) Time-dependent changes in α1 and α2 integrin expression after 24 h, 3 d, and 1 week of culture on TCPS or PEG-Scl2 hydrogels quantified by flow cytometry. Data represents six biological donors, with each data point corresponding to the average of three technical replicates per donor. C) Representative immunofluorescence images of the focal adhesion protein vinculin after 1 week of culture (n = 3 specimens per donor, 8 images per specimen). Vinculin was stained using an anti-vinculin antibody with FITC to assess focal adhesion formation. Data are presented as mean ± standard deviation across all donors (* = p < 0.05; ** = p < 0.01; ns = not significant).

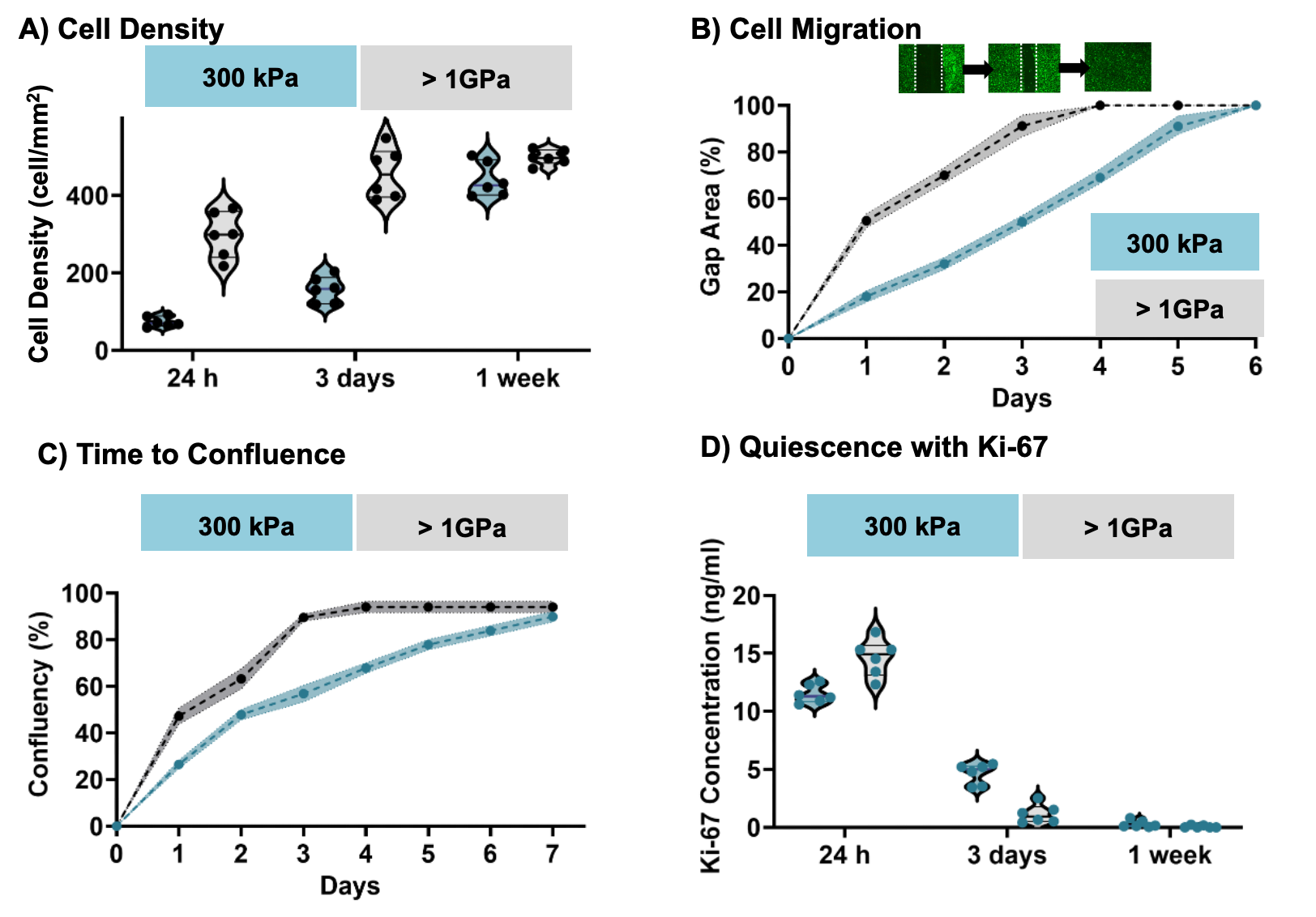

**Figure S3:** Effect of soft hydrogel expansion to stiff condition reseeding (1 GPa TCPS or 300 kPa hydrogel) on endothelial cell density, migration, confluence, and quiescence. A) Endothelial cell density quantified over time on PEG-Scl2 hydrogels (6 donors, n = 3 specimens per donor, 3 images per specimen). B) Cell migration assessed as a function of expansion condition and culture duration (6 donors, n = 3 images per timepoint). C) Time to confluence of endothelial cells (6 donors, n = 3 images per timepoint). D) Ki-67 expression used to evaluate proliferative and quiescent cell phenotypes across substrate expansion conditions (6 donors, n = 3 specimens per donor). Data represent six biological donors, with each data point corresponding to the average of three technical replicates per donor. Data for all comparisons represents the combined average of each biological and experimental replicate per donor (ns = not significant).

**Table S1:** Stiffness effect on integrin subunit expression over time.

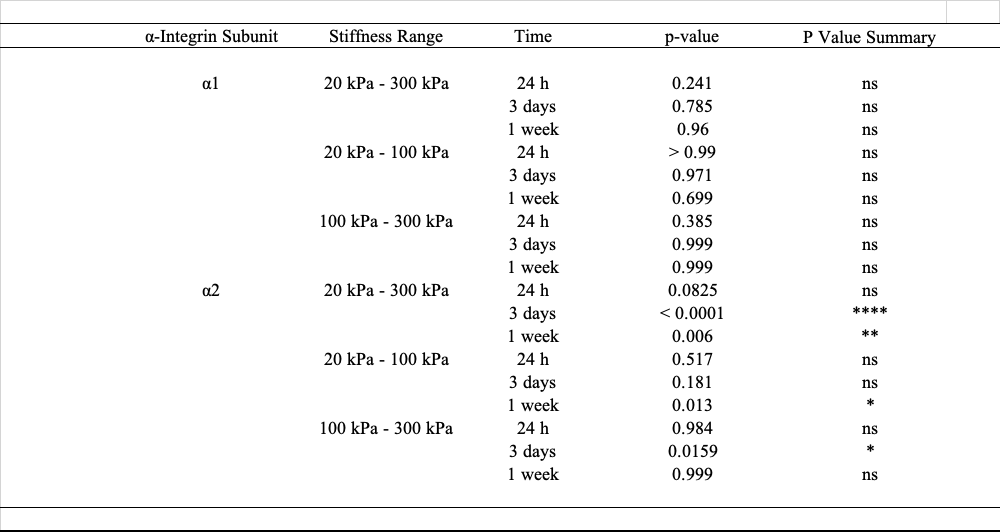

**Table S2:** Effects of expansion condition on integrin subunit expression over time.

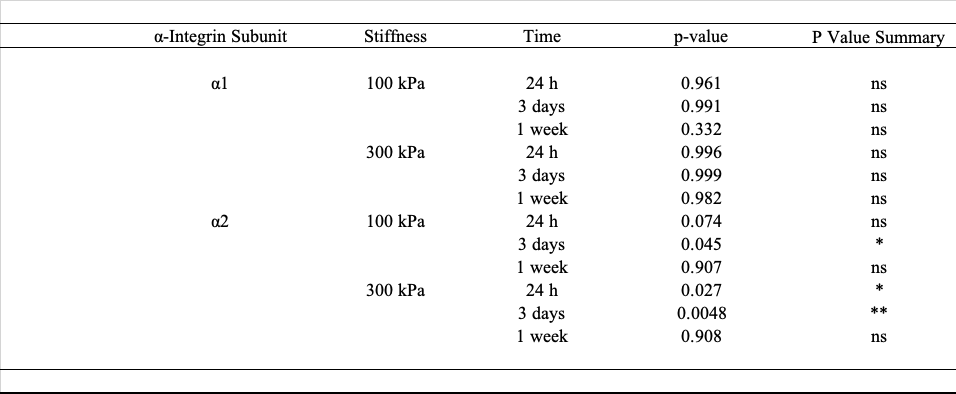
